# Epigenetic clocks and nonlinear aging in *Odocoileus* deer

**DOI:** 10.64898/2026.10.23.761432

**Authors:** Luca Montana, Cheyenne Stewart, L. Embere Hall, Aaron B. A. Shafer

## Abstract

Population age structure has important consequences for population dynamics and is a key parameter for wildlife demography and management. Epigenetic clocks have emerged as accurate molecular tools for estimating chronological age. Ungulates are among the most intensively managed wildlife worldwide, yet epigenetic clocks have been developed for only two cervid species. Here, we develop a novel epigenetic clock for mule deer (*Odocoileus hemionus*) and a multi-species clock for the *Odocoileus* genus, and use them to investigate variation in age-associated DNA methylation across the mule deer lifespan. The mule deer clock achieved accuracy comparable to recently developed mammalian epigenetic clocks (r ≈ 0.9; MAE ≈ 5% of species lifespan). The multi-species *Odocoileus* clock showed lower predictive performance (r = 0.72; MA = 0.782 years), but its broader taxonomic applicability may be particularly valuable where *Odocoileus* species occur sympatrically and hybridize. Cross-sectional epigenetic age estimates supported nonlinear age-associated methylation dynamics across the mule deer lifespan, whereas longitudinal changes in epigenetic age indicated that clocks optimized for cross-sectional age estimation may lack sufficient resolution to quantify short-term within-individual changes. Overall, our results highlight the value of integrating clock-level, CpG-level, and longitudinal analyses to investigate aging in wild populations. We also identify strongly age-associated loci that provide candidates for developing targeted, scalable, and lower-cost assays.

## Introduction

Age plays a fundamental role in shaping life histories and population dynamics and is a key parameter for wildlife management and conservation. Age dictates and corresponds to the expression of life-history traits: the onset of sexual maturity and senescence, reproductive lifespan, and age-specific patterns of survival and fecundity are inherently linked to age (Charlesworth, 1994; Monaghan et al., 2008; Stearns, 1992). When survival and reproductive success vary with age, population age structure can influence population growth through its effects on overall survival and recruitment (Hoy et al., 2020; Jackson et al., 2020). In such contexts, failure to account for age structure can create the false impression of density dependence in adult survival (Festa-Bianchet et al., 2003). For species hunted over much of their distribution, hunting can alter the age structure of populations, especially among males (Eichenberger et al., 2026; Milner et al., 2007; Schindler et al., 2017) and affect age-specific vital rates (Rughetti, 2016), making age a sought-after parameter for demographers and wildlife managers.

Despite its importance, accurate age estimates remain elusive in many wildlife monitoring projects. The most accurate age estimates typically come from studies of animal populations where individuals are monitored from birth to death. While accurate and valuable for studying individual differences in, among other traits, health and mortality risk at old ages (Gaillard & Lemaître, 2020), these long-term individual-based studies are rare, difficult to maintain, and biased toward study populations in the Northern Hemisphere (Blumstein, 2025; Clutton-Brock & Sheldon, 2010). For field studies and wildlife surveys without access to long-term data, age estimation has traditionally relied on physical features that change predictably with age. Examples include counting horn annuli in bovids (Brambilla & Canedoli, 2013; Côté et al., 1998), fish scales or growth rings in otoliths for fishes (Stevenson & Campana, 1992), tooth cementum annuli in many terrestrial mammals (Thomas, 1977), and tooth replacement-and-wear in deer (Foley et al., 2022). These methods can be accurate for certain species (e.g., tooth cementum annuli in black bears; Harshyne et al., 1998), but overall their accuracy declines in older individuals and some require experienced observers (Foley et al., 2022; Hinton et al., 2023; Rolandsen et al., 2008). Methods such as tooth wear and cementum annuli can also be influenced by environmental conditions (Foley et al., 2022; Rolandsen et al., 2008). Molecular biomarkers have emerged as less invasive alternatives for age estimation. These methods exploit biomarkers that change predictably with chronological age (i.e., the time lived by an individual since birth). Among molecular biomarkers, DNA methylation (DNAm) has proven particularly promising for chronological age estimation across animal taxa (De Paoli-Iseppi et al., 2017; Jarman et al., 2015; Le Clercq et al., 2023).

DNAm is an epigenetic modification, frequently associated to heritable or persistent changes on gene expression that occur without modifying the underlying DNA sequence (Bird, 2007). DNAm occurs at cytosines that are typically part of cytosine-phosphate-guanine dinucleotides, referred to as CpG sites. Although CpGs are relatively depleted across most of the genome, they can occur (i.e., CpG islands), often overlapping gene promoters. DNAm involves the addition of a methyl group to cytosine, generating 5-methylcytosine (Schübeler, 2015). DNAm is dynamic and can be both gained and lost with age, and some CpGs undergo predictable changes with chronological age (Field et al., 2018; Jones et al., 2015). This predictability is leveraged by epigenetic clocks to predict chronological age and was first applied across human tissues in the human pan-tissue epigenetic clock (Horvath, 2013). Animal clocks followed shortly after with the development of an humpback whale (*Megaptera novaeangliae*) epigenetic clock (Polanowski et al., 2014). Since then, species-specific epigenetic clocks have been developed across a broad diversity of animal taxa and tissue types (e.g. Haller et al., 2025; Hanski et al., 2024; Lemaître et al., 2022), along with the universal mammalian clocks (Lu et al., 2023) and a growing list of multi-species clocks (Robeck et al., 2023; Wilkinson et al., 2021; Zoller et al., 2025). Beyond chronological age estimation, DNAm can provide information on the physiological, environmental, and social factors affecting biological aging, loosely defined as a measure of tissue and organismal functional decline (Anderson et al., 2021; Cossette et al., 2023; Ryan et al., 2024). Age-associated DNAm changes may also vary across the lifespan, occurring more rapidly during juvenile growth than in later life stages (Horvath, 2013; Lemaître et al., 2022; Olecka et al., 2024; Snir et al., 2019)

Here, we investigated whether age-associated DNAm can simultaneously provide an accurate estimator of chronological age and reveal variation in the biological pace of aging across the lifespan of *Odocoileus* deer. We first developed two epigenetic clocks: a species-specific clock for mule deer (*O. hemionus*) and a multi-species *Odocoileus* clock based on mule deer and white-tailed deer (*O. virginianus*). To date, epigenetic clocks among *Cervidae* have been developed for only two wild species: white-tailed deer (Czajka et al., 2024) and roe deer (*Capreolus capreolus*; Lemaître et al., 2022). Both *Odocoileus* species are widely harvested game species of substantial ecological, social, and economic importance in North America (McShea, 2012; Mule Deer Working Group, 2026), and can hybridize (Russell et al., 2021). Accurate molecular age estimates would improve monitoring population age structure and age-specific processes. Second, we tested the hypothesis that age-associated epigenetic change is non-linear across the mule deer lifespan. We expected epigenetic age to increase more rapidly during juvenile growth than during adulthood, consistent with patterns reported in humans (Horvath, 2013; Snir et al., 2019) and roe deer (Lemaître et al., 2022). We also expected that a second shift in methylation dynamics could occur near the onset of senescence (Olecka et al., 2024). We evaluated these predictions using both cross-sectional epigenetic-clock estimates and longitudinal changes in DNAm age among repeatedly sampled individuals.

## Materials and Methods

### Study populations, sample collection and chronological age estimation

Mule deer were sampled in Wyoming as part of two monitoring projects following distinct populations: the Upper Powder River (UPR; 44° 4′N, 106° 47′W) and Wyoming Range (WYR; 42°25′00″N, 110°42′00″W) herds. Both populations are part of monitoring efforts aimed at understanding mule deer demography in Wyoming, where most herds remain below population objectives (Mule Deer Working Group, 2026). Adult females have been captured during December 2018-2021 via helicopter net-gunning (Wagler et al., 2025), and whole blood samples were collected during handling. Blood was collected from 87 individuals at initial capture, 41 of which were sampled again 364-1,099 days later (128 samples in total). Chronological age was unknown at first capture but was subsequently estimated from cementum annuli of incisors collected following mortalities and back-calculated to the date of blood sampling. WYR deer were sampled as part of a longitudinal study of individually marked animals monitored since 2013 (LaSharr et al., 2023). Nine individuals were available for this study, six of which were sampled twice 90-93 days apart (15 samples in total). Ages at time of re-capture, including blood collection, were known because the individuals were originally captured as fawns. In total, 143 mule deer samples were available for this study.

White-tailed deer were sampled in Ontario, Canada, and Texas, USA (N_Ontario_ = 26, N_Texas_ = 21) and included both sexes. Tissue samples included skin and muscles collected by local manager or provided voluntarily by hunters. Samples were stored in ethanol at the time of collection. Chronological ages were estimated using cementum annuli and tooth replacement-and-wear. Additional details on the white-tailed deer samples are provided in (Czajka et al., 2024).

### DNA extraction and methylation data

Mule deer blood samples were sent to The Clock Foundation (Torrence, CA), where DNA was extracted from whole blood. The DNA extraction for white-tailed deer is detailed in Czajka et al. (2024). Briefly, genomic DNA was extracted using the QIAGEN DNeasy Blood & Tissue kit (Qiagen, Valencia, CA), quantified using a QUBIT 3 fluorometer (Thermo Fisher Scientific), standardized to 300ng per 20 μl, and plated in 96-well plates. DNA was subjected to bisulfite conversion and profiled using the custom Illumina *HorvathMammalMethylChip40*, which targets 37,492 CpGs (Arneson et al., 2022). Arrays were processed on an iScan system by the Epigenetic Clock Development Foundation (Los Angeles, USA). Raw away data were processed in R 4.6.0 (R Core Team, 2026) using *minfi* 1.54.1 pipeline (Aryee et al., 2014; Fortin et al., 2017), and methylation levels were expressed as β values ranging from 0 (unmethylated) to 1 (fully methylated). Sample quality was assessed using *minfi*’s *detectionP* function (*P* threshold = 0.1).

### Pre-processing and final methylation datasets

Preprocessing and quality control followed Newediuk et al. (2025) and were performed separately for (i) the mule deer dataset and (ii) the combined mule deer–white-tailed deer (*Odocoileus*) dataset. Because species were assayed in different batches, batch effects in the *Odocoileus* dataset were corrected using *ComBat* in *sva* 3.56.0 (Leek et al., 2012). Approximately 0.001% of *ComBat*-corrected β values fell outside the biologically valid 0-1 range, and were truncated to 10^-7^ or 0.9999999, respectively. The *HorvathMammalMethylChip40* manifest (37,492 CpGs; https://github.com/shorvath/MammalianMethylationConsortium) was mapped to the chromosome-level mule deer and white-tailed deer assemblies (GCA_020976825.1 and GCA_023699985.4, respectively). Probes mapping to multiple genomic regions or lacking a CpG at the target position were removed. Finally, CpGs with mean DNA methylation near 0.5 were removed using the *selectProbes.middleFilter* in *MammalMethylClock* 1.0.0 (Zoller & Horvath, 2024).

Samples with compromised methylation profiles were identified using unsupervised hierarchical clustering following Zoller & Horvath (2024). Dendrogram branches were defined using *cutreeStatic* in WGCNA 1.73 (Langfelder & Horvath, 2012) with a height threshold of 0.1. To avoid pseudo-replication, only one sample per individual was used for clock training. Finally, samples were classified based on the confidence in their age estimated, and samples below the threshold value were not used to develop the epigenetic clocks.

### Epigenetic clocks validation and development

Candidate mule deer and *Odocoileus* clocks were evaluated using leave-one-out cross-validation (LOO-CV) with chronological age modeled using three transformations: identity, logarithmic, and square root. LOO-CV maximized the training sample size at each iteration and is commonly used for wildlife epigenetic clocks with limited sample sizes (Newediuk et al., 2025). Elastic-net models were fitted with α = 0.5 using *glmnet* 4.1-10 (Friedman et al., 2010). The regularization parameter λ was selected by 10-fold CV as either lambda.min, which minimized CV error, or lambda.1se, the most regularized model within one standard error of the minimum. Together, the three age transformations, two λ criteria, and two clock types produced 12 candidate models. LOO-CV was performed using *saveLOOEstimation* in *MammalMethylClock* (Zoller & Horvath, 2024), with a modified version of the function used for lambda.min models. Predictive accuracy was evaluated using median absolute error (MAE) and Pearson’s correlation coefficient (*r*) between chronological and predicted DNAm age. Accuracy was also evaluated across four defined age classes: subadults (≤ 2 years), young adults (> 2–4 years), prime-age adults (> 4–7 years), and senescent adults (> 7 years).

The transformation and λ criterion providing the best combination of MAE and *r* were used to fit each final clock to its complete training dataset. Final models were fitted by elastic-net regression (α = 0.5) using lambda.min from 10-fold *cv.glmnet*, which selected the CpGs and corresponding regression coefficients retained in each clock. Finally, the performance of the mule deer and *Odocoileus* clocks was compared with that of the published universal mammalian (Lu et al., 2023), white-tailed (Czajka et al., 2024), and roe deer (Lemaître et al., 2022) clocks. Published clock coefficients were applied to normalized, non-batch-corrected β values for mule deer and white-tailed deer, and predicted DNAm ages were back-transformed according to the transformation used by each clock. Performance was compared using MAE and *r*.

### Age-associated methylation dynamics

We used three complementary approaches to assess whether age-associated epigenetic change varied across the mule deer lifespan: cross-sectional DNAm-age predictions, longitudinal changes in DNAm age, and CpG-specific methylation trajectories. First, DNAm ages obtained by LOO-CV were regressed against untransformed chronological age using three models: (i) a linear model, (ii) a quadratic model, and (iii) a generalized additive model (GAM) with a cubic regression spline (*mgcv* 1.9-4; Wood, 2011). Model support was compared using corrected Akaike Information Criterion (AICc) and Akaike weights (Burnham et al., 2011).

Second, the final mule deer clock was applied to the 44 individuals of unknown age sampled twice. For each individual, Δ_age_ was calculated as the difference in predicted DNAm age between the second (*t_1_*) and first (*t_0_*) sampling events. Individuals were assigned to the four age classes defined above based on DNAm age at *t_0_*, and differences in Δ_age_ among classes were tested using a Kruskal–Wallis test followed by pairwise Mann–Whitney *U* tests in R.

Finally, an epigenome-wide association study (EWAS) was used to identify CpGs associated with chronological age and characterize their age-associated methylation trajectories. The CpGs retained after preprocessing were analyzed using *lmFit* in *limma* 3.68.4 (Ritchie et al., 2015), with normalized β values modeled as a function of chronological age and study population (UPR or WYR). Significant associations were identified using both a Bonferroni-corrected P < 0.05 threshold and a Benjamini–Hochberg false discovery rate (FDR) < 0.05 (Benjamini & Hochberg, 1995; Huang et al., 2020). Significant CpGs were subsequently annotated to genomic features and nearby genes. Genomic annotations were used to test whether (1) age-associated CpGs were enriched or depleted in particular genomic regions, (2) the direction of age-associated methylation differed among genomic regions, and (3) significantly age-associated CpGs within each region are disproportionally hyper- or hyo-methylated relative to non-significant CpGs. For (1), a *χ*^2^ goodness-of-fit test was used to determine whether the distribution of significant CpGs among genomic regions differed from that expected based on the distribution of all CpGs included in the EWAS (N = 27,270); region-specific enrichment or depletion was subsequently assessed using Fisher’s exact test, with odds ratios (OR) quantifying effect sizes. For (2), a *χ*^2^ test of independence was used to determine whether the proportions of significantly hyper- and hypo-methylated CpGs differed among genomic regions. For (3), Fisher’s exact test on 2-by-2 contingency tables were used within each genomic region to test the association between methylation direction (hyper-vs hypo-methylation) and EWAS significance (significant vs non-significant), with odds ratios quantifying the direction and magnitude of enrichment. For both sets of region-specific Fisher’s exact tests (analyses 1 and 3), P-values were adjusted for multiple testing using the Benjamini–Hochberg procedure (Benjamini & Hochberg, 1995). All statistical analyses were performed in R.

## Results

### Mule deer and Odocoileus epigenetic clocks

A total of 50 individuals ranging from 1.5 to 12 years old passed the pre-processing filtering along with 27,270 CpGs to train the mule deer epigenetic clocks. According to the performance of the LOO-CV clocks, models using the λ = lambda.min yielded the most accurate results compared with those using λ = lambda.1se (**Figure S1**), and the square-root-transformed age LOO-CV models provided the best combination of MAE and *r* (MAE = 0.636 years, *r* = 0.88; **Figures 1a** and **S1**). The accuracy of the square-root-transformed age LOO-CV clocks appeared to decrease with for older individuals (MAE_subadults_ = 0.345, MAE_young adults_ = 0.471, MAE_prime age_ = 0.678, MAE_senescent_ = 1.685). The final elastic-net model (α = 0.5, λ = 0.012, square-root-transformed chronological age) selected 65 CpGs (**Figure S2**) which were used to develop the mule deer epigenetic clock (**Figure 1b**).

**Figure 1.**
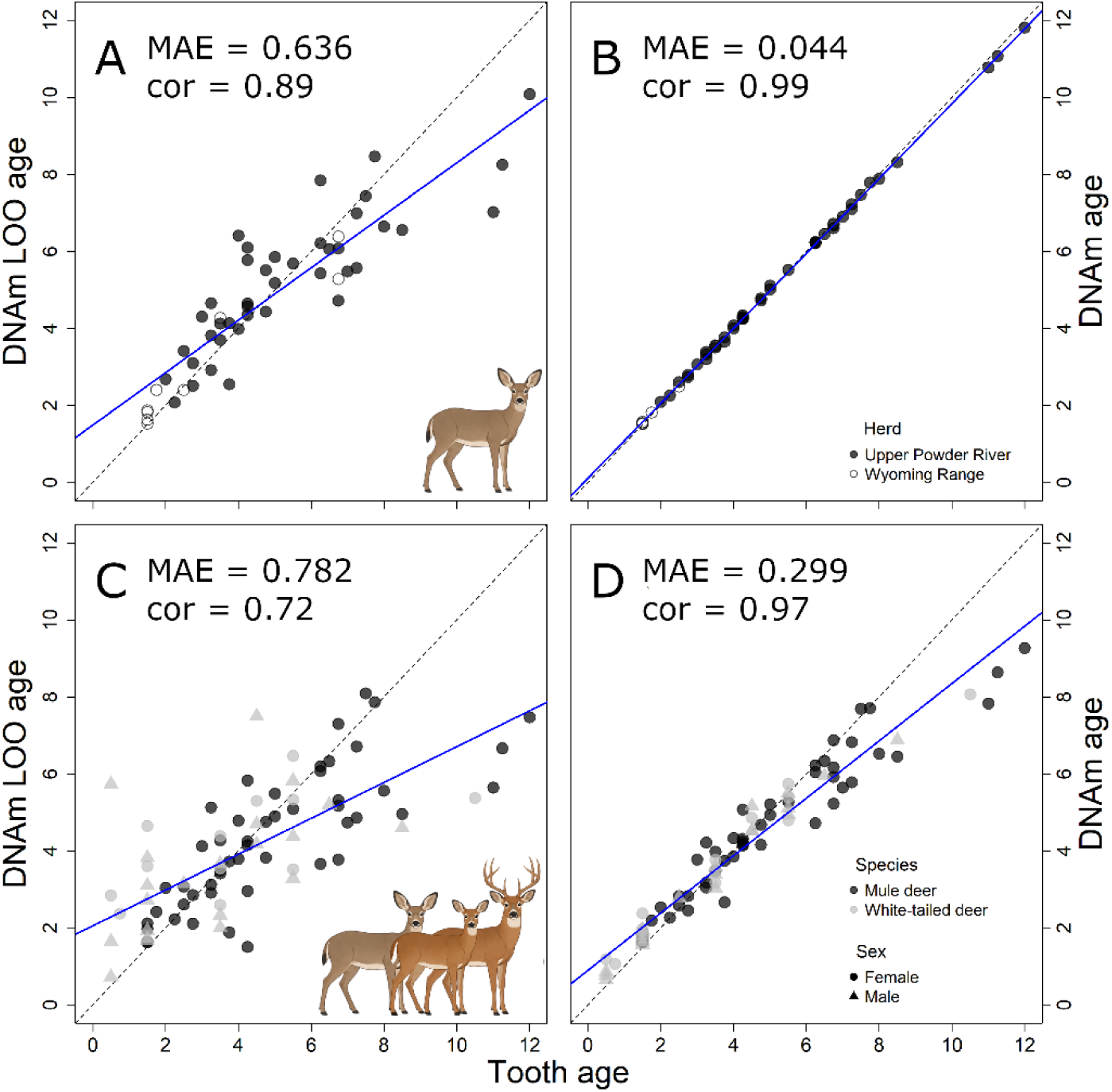
Leave-one-out cross-validation (LOOO-CV; **A**, **C**) and final epigenetic clocks (**B**, **D**) showing the relationship between chronological (tooth) age and DNA methylation age predictions (expressed in years) for mule deer (N = 50; upper row) and the *Odocoileus* genus (N = 83; bottom row). The mule deer LOO-CV (**A**) and final (**B**) clocks were trained using the λ value minimizing the CV error and with square-root-transformed chronological ages. The Odocoileus clocks (**C**-**D**) were also trained using the λ value minimizing the CV error but with log-transformed chronological ages. Solid blue lines represent the linear regression of epigenetic age, and the dashed line is the diagonal (x = y). Predictive accuracies for the LOO-CV and final clocks (median absolute error [MAE] and Pearson’s correlation [cor]) are reported in each panel. The deer icons were made on ChatGPT.

The *Odocoileus* epigenetic clocks were trained using the 50 mule deer included in this study together with the 33 white-tailed deer ranging from 0.5 to 10.5 years used by Czajka et al. (2024) to develop the white-tailed deer epigenetic clock (N_tot_ = 83) and 27,112 CpGs. As for the *Odocoileus* clock, LOO-CV models trained with λ = lambda.min yielded, on average, the most accurate results, whereas the log-transformed LOO-CV models showed the best combination of MAE and *r* (MAE = 0.782, *r* = 0.72, **Figure 1c** and **S3**). The *Odocoileus* clock was therefore developed using log-transformed chronological ages and 66 CpGs (α = 0.5, λ = 0.075; **Figures 1d** and **S4**).

Compared with the mule deer clock, the *Odocoileus* clock estimated mule deer ages less accurately (*r*_mule_ = 0.89 and MAE_mule_ = 0.636 versus *r_Odocoileus_* = 0.72 and MAE*_Odocoileus_* =0.782 years). However, the *Odocoileus* clock was able to estimate the age of animals of both species, whereas neither the mule deer clock nor the white-tailed deer clock (Czajka et al., 2024) were able to accurately estimate the age of individuals of the other species (**Figure S5**). The roe deer clock (Lemaître et al., 2022) also failed to accurately estimate the age in either species (**Figure S5**). In contrast, the universal mammalian clocks 1 and 2 (Lu et al., 2023) performed well for both species (MAE_clock1_ = 0.880, *r*_clock1_ = 0.793; MAE_clock2_ = 0.732, *r*_clock2_ = 0.825; **Figure S5**). Both universal mammalian clocks showed stronger correlations between the DNAm age and chronological age than the *Odocoileus* clock, but only the universal mammalian clock 2 achieved a lower MAE than the *Odocoileus* clock.

### Aging dynamics of mule deer and EWAS of chronological age

To test whether aging was constant across all life stages of mule deer or whether a non-linear relationship was present between DNAm age and chronological age, we used the DNAm ages estimated from the square-root--transformed LOO-CV models as the response variable for the three regression models (linear model, quadratic model, cubic spline GAM model; **Table 1a**, **Figure 2**). Within the candidate set, the two non-linear models received substantially greater support (ΔAICc < 1.05) than the linear model (ΔAICc = 6.41), together accounting for roughly 97.5% of the Akaike weight and receiving approximately 39 times greater support than the linear model. The model selection provided little evidence for distinguishing the precise form of non-linearity (**Table 1b**).

**Figure 2.**
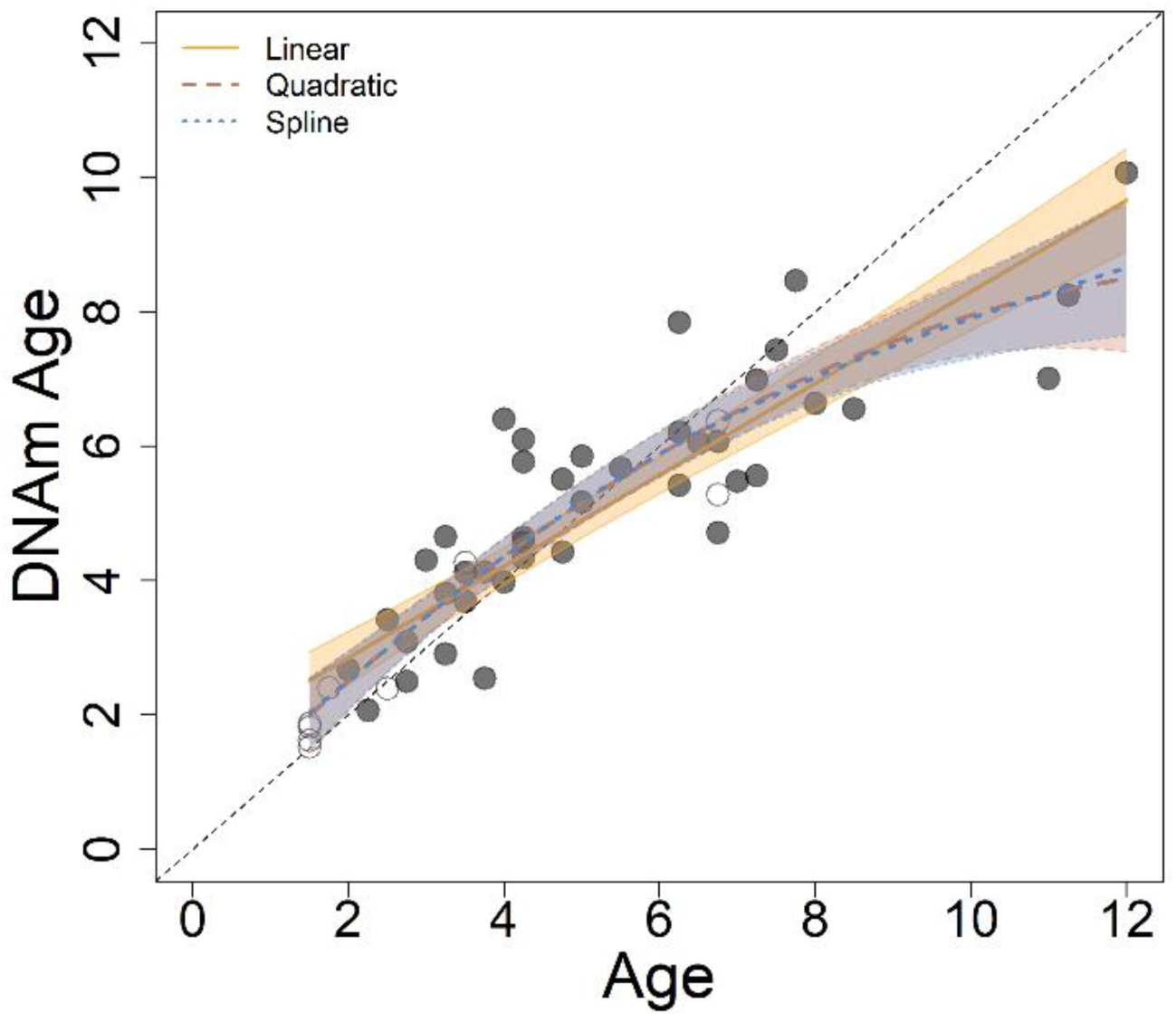
Relationship between predicted DNAm age (expressed in years) estimated from the leave-one-out cross-validation (LOO-CV) clocks (Figure 1A) and untransformed chronological age for mule deer sampled in Wyoming, 2018-2022. Mule deer originated from two sample sites: Upper Powder River (black dots) and Wyoming Range (black circles). Lines and associated shaded areas represent the linear and non-linear models used to study the relationship.

**Table 1.**
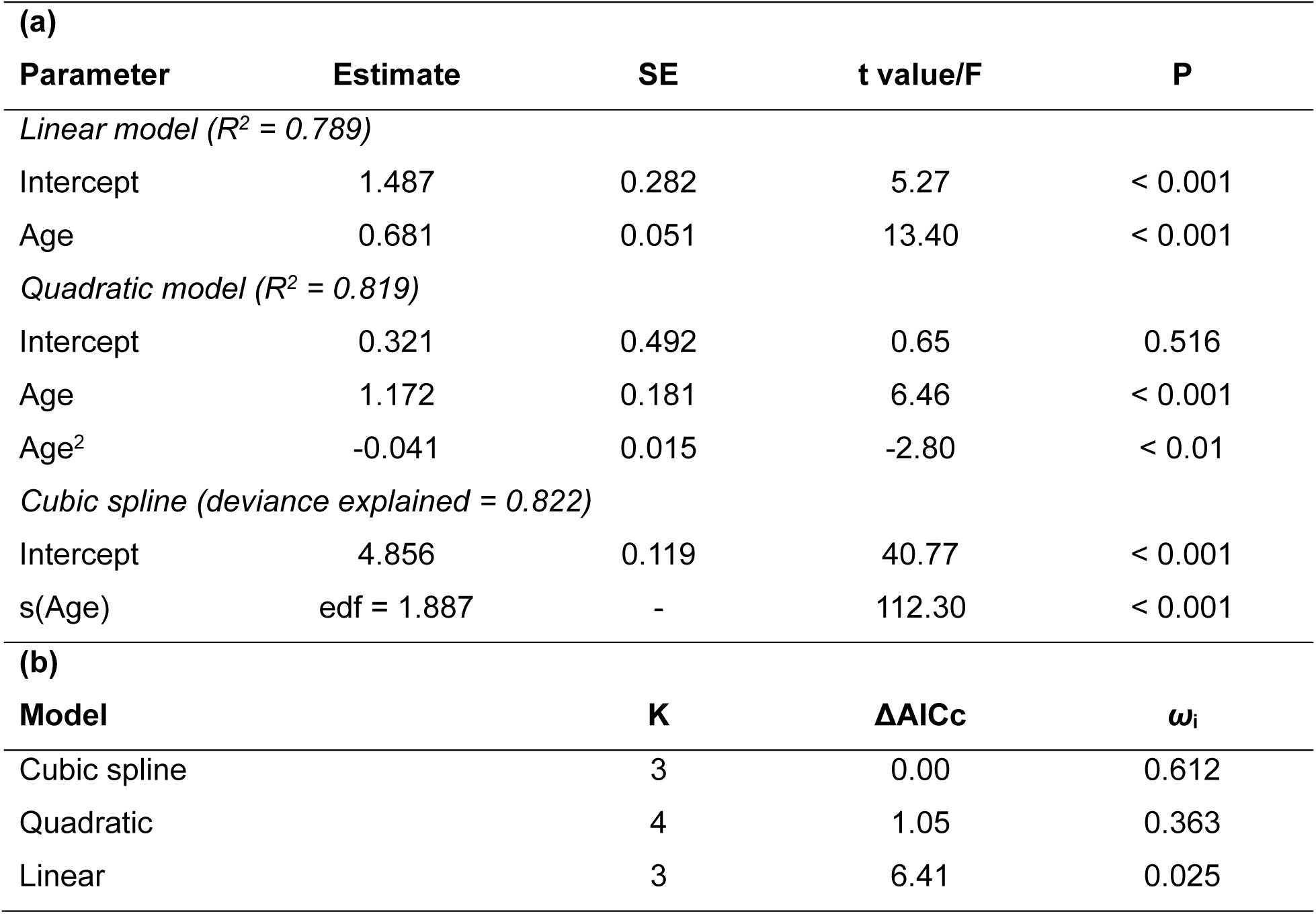
Estimates of linear, quadratic and cubic spline effects of chronological age on DNAm age (a) and model selection aiming to quantify if the relationship between chronological and DNAm age was linear or non-linear (b) in mule deer samples in Wyoming, 2018-2022. (a) For each regression model, the estimates, associated standard error (SE), t-value (or F value for the cubic spline), are provided along with the proportion of variance/deviance explained by each model. (b) For each candidate model, the number of estimated parameters (K), the difference in AIC_c_ value (ΔAIC_c_) and Akaike weight (*ω_i_*) are provided.

| **(a)** |  |  |  |  |
| --- | --- | --- | --- | --- |
| **Parameter** | **Estimate** | **SE** | **t value/F** | **P** |
| *Linear model ($R^2 = 0.789$)* |  |  |  |  |
| Intercept | 1.487 | 0.282 | 5.27 | < 0.001 |
| Age | 0.681 | 0.051 | 13.40 | < 0.001 |
| *Quadratic model ($R^2 = 0.819$)* |  |  |  |  |
| Intercept | 0.321 | 0.492 | 0.65 | 0.516 |
| Age | 1.172 | 0.181 | 6.46 | < 0.001 |
| Age<sup>2</sup> | -0.041 | 0.015 | -2.80 | < 0.01 |
| *Cubic spline (deviance explained = 0.822)* |  |  |  |  |
| Intercept | 4.856 | 0.119 | 40.77 | < 0.001 |
| s(Age) | edf = 1.887 | - | 112.30 | < 0.001 |
| **(b)** |  |  |  |  |
| **Model** | **K** |  | **$\Delta\text{AIC}_c$** | **$\omega_i$** |
| Cubic spline | 3 |  | 0.00 | 0.612 |
| Quadratic | 4 |  | 1.05 | 0.363 |
| Linear | 3 |  | 6.41 | 0.025 |

When applying the mule deer clock developed here to individuals re-sampled in the field (time elapsed between sampling events = 90-1,099 days), samples at *t*_1_ were, on average, older than at *t*_0_. (**Figure S6**), although deer sampled after about 365 days were often predicted to be younger at *t*_1_ compared to *t*_0_ (**Figure S6**). We then investigated whether these estimates were influenced by the age of the deer at *t*_0_. To do this, we subdivided the animals into age classes (N_subadult_ = 3, N_young adults_ = 12, N_prime age adults_ = 23, N_senescent_ = 6) and examined whether any age class systematically underestimated DNAm age relative to the time elapsed between sampling events. The average Δ_age_ was 108.93 days for subadults, 407.4 days for young adults, 372.5 days for prime-age adults, and 92.4 days for senescent adults (**Figure 3**). Despite the apparent differences in mean Δ_age_, the comparisons among age classes were not significant (Mann-Whitney *U* tests: subadults v young adults: *U* 8, *P* = 0.180, young adults v prime-age adults: *U* 165, *P* = 0.362, prime-age adults v senescent adults: *U* 47, *P* = 0.336; Kruskal-Wallis test including all age classes: *H* = 2.404, df = 3, *P* = 0.493; Kruskal-Wallis test excluding subadults: *H* = 1.616, df = 1, *P* = 0.446).

**Figure 3.**
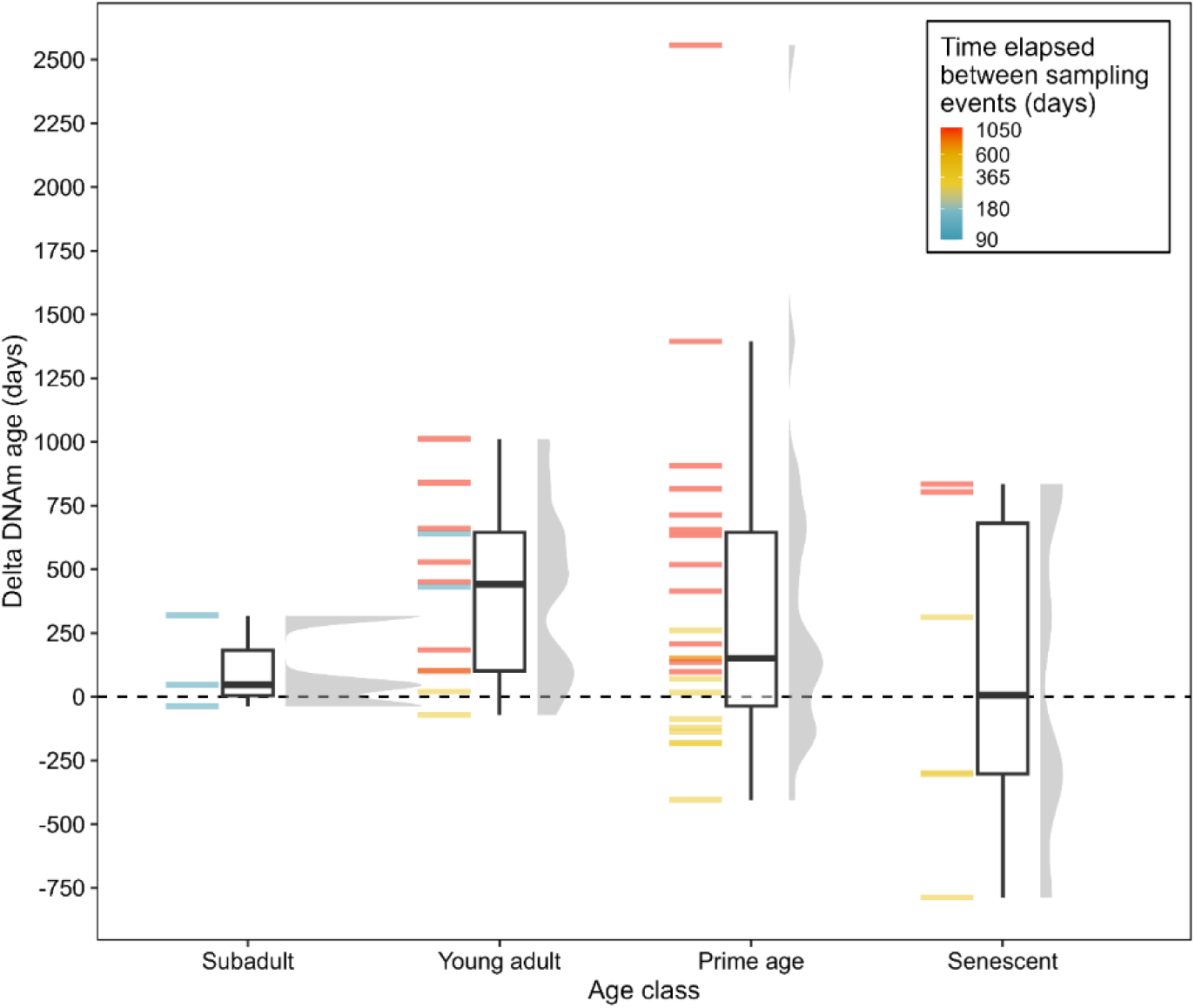
Comparison of change in DNAm age (Δ_age_) among 44 mule deer from different age classes (subadults: ≤ 2 years; young adults: > 2–4 years; prime-age adults: > 4–7 years; senescent adults: > 7 years) sampled in Wyoming, 2018-2022. For each age class, the distribution of Δ_age_ is shown using a half-eye plot: the boxplot indicates the median and interquartile range, the grey half-violin indicates the density distribution. Individual observations are represented by horizontal marks, colored according to the time elapsed between sampling events.

The EWAS revealed that chronological age was associated with methylation at a substantial proportion of the 27,270 CpGs of the methylation array that mapped on both mule deer and white-tailed deer genome assemblies (CpGs significantly correlated with age identified by Bonferroni *P* correction: 40 CpGs or about 0.15% of CpGs; **Figure 4a**). Among these 40 CpGs, 27% showed decreasing methylation levels with age (**Figure S7**). We did not observe complete (de)saturation of methylation levels at any site (**Figures 4c**, **S7**). However, in several cases the increase or decrease of methylation appeared to taper off at older ages (e.g., both AMER2 CpGs, LHFPL4, INSM1, both CDX2 CpGs, FOXP4, POU4F1, PAX5, ZNF536). The hypo- and hypermethylation patterns of CpGs identified using the Bonferroni corrected significance threshold or by the Benjamini–Hochberg FDR were similar (**Figure 4c**), despite the latter identified more CpGs associated with age (N = 1,733 CpGs). Because few CpGs met the Bonferroni-corrected significance threshold, analyses of genomic region enrichment and differences in hyper- and hypomethylation across regions were restricted to FDR-significant CpGs. The genomic distribution of age-associated CpGs differed from that expected based on the distribution of all CpGs included in the EWAS (χ² = 262.96, df = 5, P < 0.001). Specifically, age-associated CpGs showed significant enrichment or depletion across most genomic regions, except for 5′ UTRs, and were disproportionately hypermethylated in promoters, 5′ UTRs, intergenic regions, and exons relative to non-significant CpGs (**Figure 4b**).

**Figure 4.**
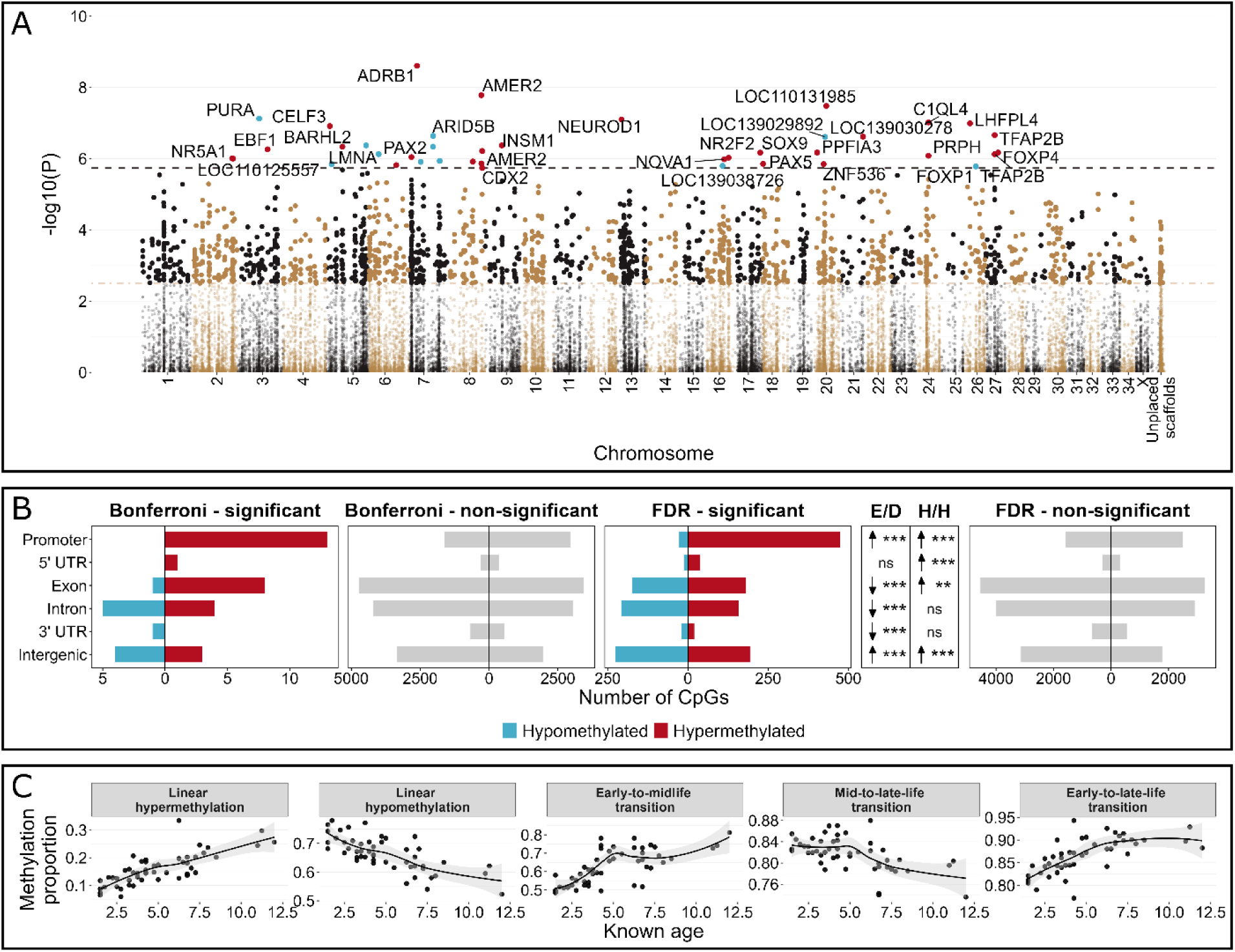
Epigenome-wide association study (EWAS) of chronological age in mule deer and typical methylation trajectories during aging. (**A**) Manhattan plot of the EWAS of chronological age. Data points represent CpG sites (N = 27,270). Genomic coordinates are based on the alignment of the CpG probes to the white-tailed deer genomic assembly (alternate colors between chromosomes). The black and golden brown horizontal dashed lines identify CpGs significantly correlated with age according to the Bonferroni-corrected *P* and the Benjamini–Hochberg False Discovery Rate (FDR) methods, respectively. The direction of associations with Bonferroni-corrected *P* < 0.05 is highlighted by red (hypermethylated) and blue (hypomethylated) colours, and CpGs are labeled by the neighboring genes. (**B**) Distribution of hyper- and hypomethylated CpGs across genomic regions, separated by significance status under Bonferroni and FDR corrections. Grey bars represent CpGs not significantly associated with age. Statistical comparisons were performed using FDR-significant CpGs only. The genomic distribution of age-associated CpGs differed significantly from the background distribution of all CpGs (χ² = 262.96, df = 5, P < 0.001), while methylation direction differed among genomic regions (χ² = 342.91, df = 5, P < 0.001). Arrows indicate significant enrichment (upward-pointing arrows) and depletion (downward-pointing arrows) of age-associated CpGs relative to the genomic background (E/D), or significant enrichment of hypermethylated CpGs relative to the directional distribution of non-significant CpGs within each region (H/H). Significance was determined using Fisher’s exact tests with Benjamini–Hochberg correction (adjusted P < 0.05). (**C**) Typical methylation trajectories identified across the CpGs significantly correlated with age according to the Bonferroni-corrected *P*.

## Discussion

This study presents a newly developed epigenetic clock for mule deer and a multi-species clock for the *Odocoileus* genus (**Figure 1**), which includes widely harvested game species of substantial ecological, social, and economic importance in North America. The accuracy of the mule deer clock is comparable to that of other recently developed mammalian epigenetic clocks (*r* ≈ 0.9, MAE ≈ 5% of the lifespan of the species of interest; e.g., Cossette et al., 2023; Hanski et al., 2024; Lemaître et al., 2022), highlighting its potential as a new standard for molecular age estimation in this species. The *Odocoileus* clock provides an alternative for age estimation across the genus, particularly because species-specific clocks developed for either mule deer or white-tailed deer perform poorly when applied to the other species (this study, Czajka et al., 2024). The taxonomic flexibility of the *Odocoileus* clock could also be particularly valuable where the two species occur sympatrically and hybridize, as it occurs repeatedly across western North America (e.g. Russell et al., 2021), although its performance in genetically confirmed hybrids remains to be tested.

This study also provides evidence that age-associated methylation dynamics are nonlinear across the mule deer lifespan, with faster DNAm change during earlier life and evidence of altered methylation trajectories again near the transition to senescence (Horvath, 2013; Lemaître et al., 2022; Snir et al., 2019). Interestingly, the majority of CpGs associated with chronological age become increasingly methylated with age, consistent with the tendency of CpG islands to become hypermethylated against a background of genome-wide age-dependent hypomethylation (Johansson et al., 2013; Jones et al., 2015). Inspection of age-associated CpGs further revealed trajectories ranging from linear hyper- or hypomethylation to non-linear trajectories characterized by transitions in methylation changes between different life stages (Olecka et al., 2024).

### Non-linear age-associated methylation across lifespan

Evidence from multiple species suggests that age-associated DNAm changes do not proceed at a constant rate across the lifespan, but instead can follow logarithmic or log-linear trajectories, with a rapid rate of epigenetic change in juveniles, followed by either progressively slower change through adulthood (logarithmic trend; Snir et al., 2019) or a more constant rate across post-developmental stages (log-linear trend; Horvath, 2013). Accordingly, chronological age is often transformed before epigenetic-clock training, allowing linear predictive models to accommodate non-linear associations between chronological age and DNAm. Consistent with such non-linearity, age transformation improved prediction in both clocks (**Figures S1**, **S3**).

Importantly, the relationship between epigenetic and untransformed chronological age in mule deer was also better described by a non-linear model despite the absence of very young individuals from our dataset (the youngest deer in our dataset was 1.5 years old, around the age of sexual maturity in female mule deer). In roe deer, the quadratic relationship between DNAm and chronological age disappeared when considering only adults (individuals > 1 year old; Lemaître et al., 2022). The persistence of non-linearity in our largely post-developmental dataset therefore suggests that nonlinear age-associated methylation in mule deer may not be driven solely by rapid juvenile development. Here, the relationship changed at approximately 6 years of age, well after female mule deer reach sexual maturity but closer to the onset of senescence, raising the possibility of an additional shift in methylation dynamics during the transition from prime adulthood to later life. Two non-mutually exclusive explanations could account for this pattern. First, the apparent late-life flattening could partly reflect reduced predictive accuracy at older ages, because senescent individuals were less represented during clock training. Such age-related prediction bias has also been reported in delphinid epigenetic clocks with limited representation of older individuals (Barratclough et al., 2024; Martien et al., 2026). Second, the pattern could reflect a genuine shift in age-associated methylation during the mid-to-late-life transition, as described in mice (Olecka et al., 2024).

The latter interpretation is supported by the methylation profiles of age-associated CpGs identified by the EWAS. About 50% presented non-linear methylation trajectories, including the early-to-mid-life and mid-to-late-life transitions described by Olecka et al. (2024), as well as an additional pattern that we term the early-to-late-life transition. This latter trajectory, detected in 14 methylation profiles from 12 CpGs (30% of CpGs), was characterized by a steady methylation increase from early life until the onset of senescence, followed by a late-life plateau despite methylation levels remaining below apparent saturation. Because these trajectories were observed directly at age-associated CpGs rather than being derived from clock predictions, they provide partially independent support for a biological change in methylation dynamics at older ages. This pattern suggests that age-associated methylation changes can differ not only in their timing but also in their duration, with some changes extending across most of the lifespan before reaching a late-life plateau.

### Longitudinal changes in DNAm age

The longitudinal dataset showed that DNAm age tended to increase between successive sampling events for most individuals, although decreases were not uncommon. The mule deer clock has an average accuracy of ± 7 months, but prediction error increased with chronological age (up to about ± 20 months for senescent deer). Consequently, short sampling intervals (< 1 year) may fall within the prediction error and thus be insufficient to detect biologically meaningful changes in DNAm age in long-lived species such as deer, as also noted for Asian elephants (*Elephas maximus*; Arai & Inoue-Murayama, 2026). This highlights an important distinction between cross-sectional age estimation and longitudinal applications of epigenetic clocks: a clock can estimate chronological age accurately across a broad lifespan while still lacking sufficient resolution to detect relatively small within-individual changes over short sampling intervals.

Nevertheless, both increases and decreases in estimated DNAm age have been reported in several wildlife studies with longitudinal sampling (Anderson et al., 2021; Lemaître et al., 2022; Yagi et al., 2024), with particularly clear examples in hibernating animals (Pinho et al., 2022; Sullivan et al., 2022). Given the magnitude of prediction error in our clock, however, decreases observed in individual mule deer should not necessarily be interpreted as biological epigenetic rejuvenation or deceleration.

Of note, the mean change in DNAm age (Δ_age_) appeared to decrease among age classes, although differences were not statistically significant. This apparent pattern should be interpreted cautiously, however, because prediction error was large relative to the expected change in DNAm age over short sampling intervals. In particular, individuals sampled approximately one year apart showed, on average little to no increase in predicted DNAm age. Because these individuals were not excluded a priori, we do not interpret their removal as a separate biological test. Rather, this sensitivity comparison showed that the apparent age-class pattern was strongly influenced by observations for which the expected temporal change in DNAm age approached the prediction error of the clock. When these observations were omitted, mean Δ_age_ was similar among young, prime-age, and senescent adults, suggesting that the longitudinal data provide little support for large differences in DNAm-age accumulation among adult age classes (Horvath, 2013; Lemaître et al., 2022). This does not necessarily contradict the cross-sectional clock predictions and CpG-level trajectories, which suggested slower age-associated epigenetic change near the onset of senescence. Instead, the discrepancy likely reflects the different temporal and biological scales captured by the three approaches: cross-sectional models and CpG trajectories characterize age-associated differences accumulated over several years, whereas the longitudinal analysis attempts to resolve comparatively small within-individual changes over much shorter intervals.

### Conclusions

The mule deer and Odocoileus epigenetic clocks developed here provide minimally invasive tools for estimating chronological age, with potential applications in wildlife monitoring and management. Beyond age estimation, our findings reveal nonlinear methylation dynamics across the mule deer lifespan, including changes extending beyond juvenile development and potentially associated with the onset of senescence. These results highlight the value of combining clock-level, CpG-level, and longitudinal approaches to investigate aging in wild populations. Further validation across populations and geographic regions could strengthen the broader applicability of these clocks. Ultimately, translating strongly age-associated CpGs into targeted, scalable, and cost-effective assays will be essential for integrating epigenetic age estimation into routine wildlife monitoring and management.

## Acknowledgements

We thank the volunteers and personnel of the Wyoming Game and Fish Department for assistance with data collection and sample processing during captures and with mortality recoveries of mule deer from the Upper Powder River herd. Kevin L. Monteith provided mule deer samples from the Wyoming Range, and Alan Cain provided white-tailed deer samples from Texas. We thank Marco Festa-Bianchet for discussions on the influence of age structure on wildlife population dynamics, and Levi Newediuk, Sarah Wolf, and members of the Wildlife Epigenetic Clock community for insightful discussions on epigenetic clocks. We also thank members of the Wildlife and Applied Genomics group at Trent University, particularly Irina Badell, Marie-Laurence Cossette, and Natalie Czajka, for helpful discussions on the bioinformatics analyses. This work was supported by a Natural Sciences and Engineering Research Council of Canada Discovery Grant (ABAS), the Digital Research Alliance of Canada for Research Group (ABAS), and the Wyoming Gam and Fish Commission (grant number: 005543).

## Author Contributions

All authors co-conceived and co-designed the data. CS, LEH, and ABAS collected and acquired field data. LM performed all bioinformatic and statistical analyses and wrote the first draft of the paper. All authors interpreted that results, as well as revised and contributed critically to the manuscript.

## Ethics Approval Statement

All applicable institutional and national guidelines for the care and use of animals were followed. Animal-handling protocols were approved by Institutional Animal Care and Use Committees [Wyoming Range, University of Wyoming #20200305KM00412; Upper Powder River, Wyoming Game and Fish Department #18-02] and authorized under Wyoming Game and Fish permits 33-985 (Wyoming Range) and 33-1197 (Upper Powder River).

## Data Availability Statement

Raw epigenetic data will be deposited on NCBI Sequence Read Archive (SRA). Associated metadata and scripts will be available on Zenodo and in a public GitHub repository.

