## Supplementary Figures for "Epigenetic clocks and nonlinear aging in *Odocoileus* deer"

Electronic Supplementary Material for

### Abstract

Population age structure has important consequences for population dynamics and is a key parameter for wildlife demography and management. Epigenetic clocks have emerged as accurate molecular tools for estimating chronological age. Ungulates are among the most intensively managed wildlife worldwide, yet epigenetic clocks have been developed for only two cervid species. Here, we develop a novel epigenetic clock for mule deer (Odocoileus hemionus) and a multi-species clock for the Odocoileus genus, and use them to investigate variation in age-associated DNA methylation across the mule deer lifespan. The mule deer clock achieved accuracy comparable to recently developed mammalian epigenetic clocks ($r \approx 0.9$; MAE ≈ 5% of species lifespan). The multi-species Odocoileus clock showed lower predictive performance ($r = 0.72$; MA = 0.782 years), but its broader taxonomic applicability may be particularly valuable where Odocoileus species occur sympatrically and hybridize. Cross-sectional epigenetic age estimates supported nonlinear age-associated methylation dynamics across the mule deer lifespan, whereas longitudinal changes in epigenetic age indicated that clocks optimized for cross-sectional age estimation may lack sufficient resolution to quantify short-term within-individual changes. Overall, our results highlight the value of integrating clock-level, CpG-level, and longitudinal analyses to investigate aging in wild populations. We also identify strongly age-associated loci that provide candidates for developing targeted, scalable, and lower-cost assays.

### *Supplementary Figures*

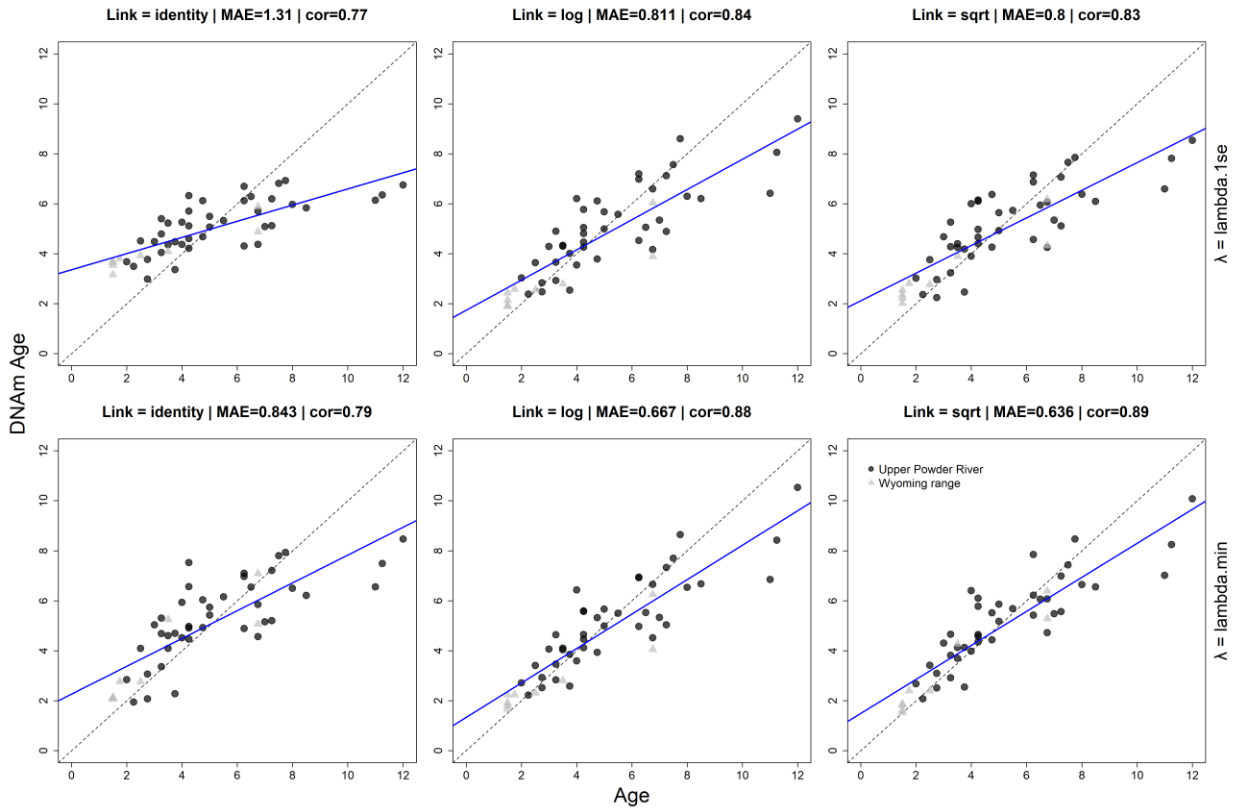


**Figure S1.** Evaluation of tooth-age transformation (first column: identity; second column: logarithmic; third column: square root) and regularization parameter λ (upper row: λ value minimizing the cross-validation [CV] error; lower row: λ corresponding to the most regularized model within one standard error of the minimum) on the predictive accuracy of leave-one-out cross-validation (LOO-CV) clocks for mule deer sampled in Wyoming, 2018-2022. Mule deer samples were collected from two different areas: Upper Powder River (N = 41) and Wyoming Range (N = 9). The fitted linear regression is represented by the blue line. Predictive accuracy of LOO-CV clocks was evaluated using the median absolute error (MAE) and Pearson's correlation coefficient (indicated above each panel). The dashed line represents the diagonal (x = y).

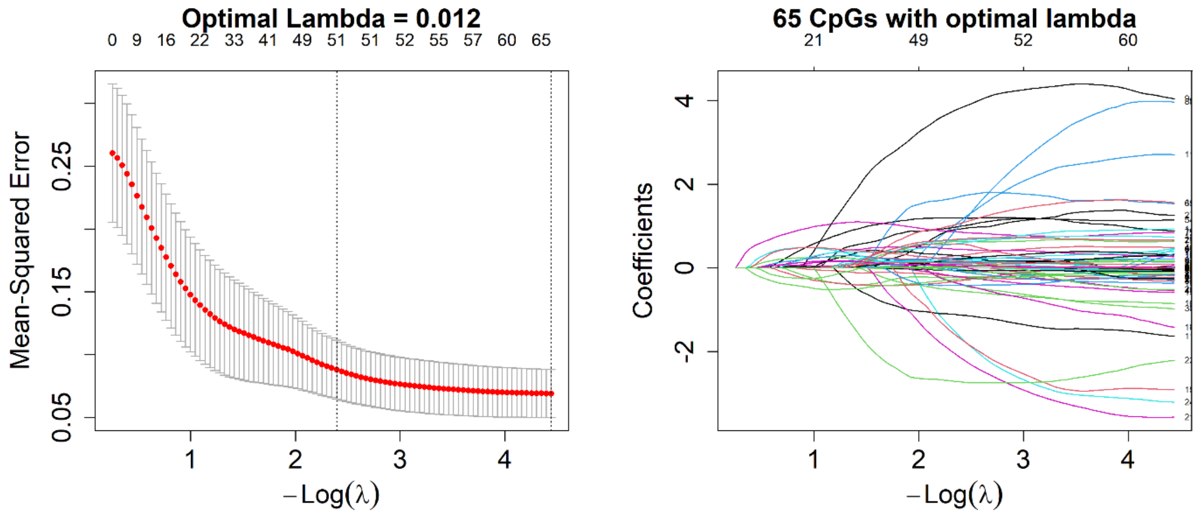


**Figure S2.** Selection of the optima regularization parameter (λ) and visualization of the regularization paths across λ values for the mule deer 10-fold cross-validated elastic-net model (α = 0.5), using square-root-transformed tooth age. Left panel: cross-validation curve used to select the optimal λ. Each point (red dot) represents the mean squared CV error, with error bars indicating ± SD. The vertical dotted lines represent the λ minimizing the CV error (lambda.min; here regarded as optimal λ here) and λ corresponding to the most regularized model within one standard error of the minimum (lambda.1se). Right panel: regularization path of CpG coefficients across λ values. Each curve corresponds to a CpG site (N = 27,270 CpGs), and CpGs with non-zero coefficients at the optimal λ were selected to fit the final mule deer elastic-net model.

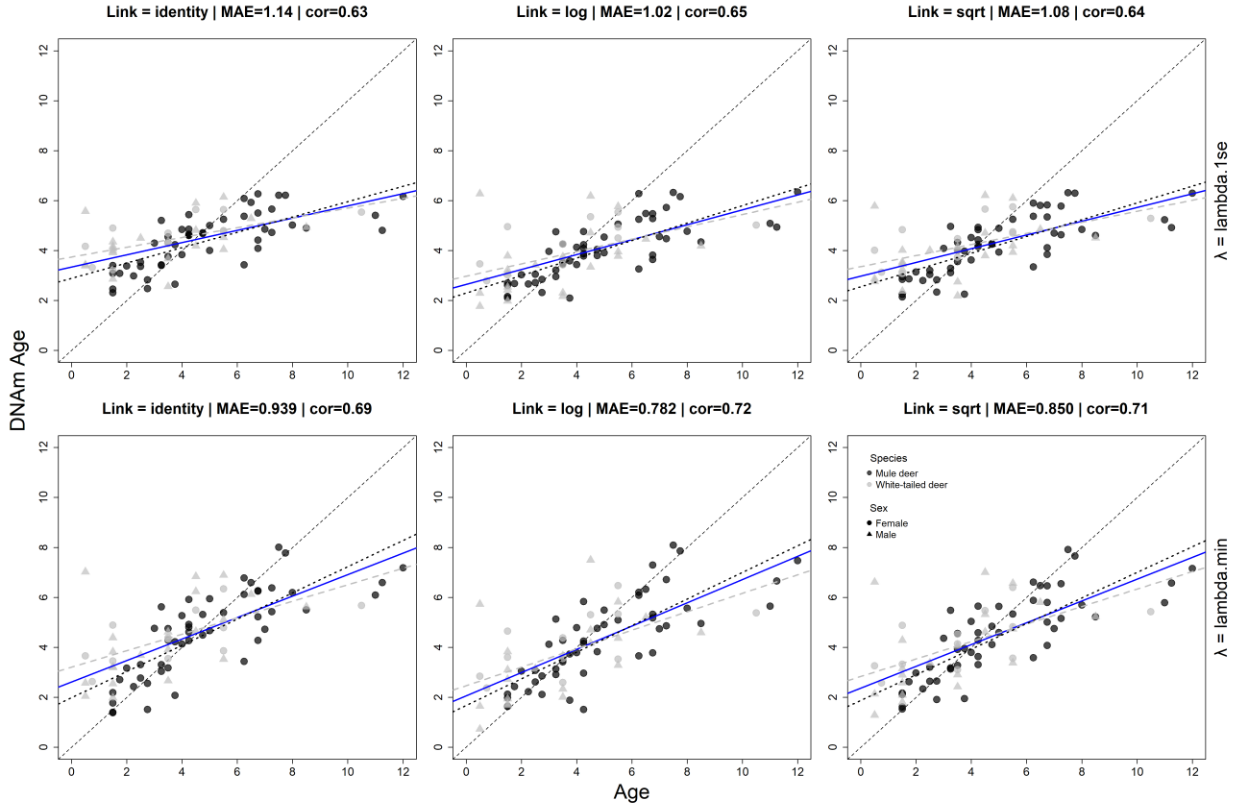


**Figure S3.** Evaluation of tooth-age transformation (first column: identity; second column: logarithmic; third column: square root) and regularization parameter λ (upper row: λ value minimizing the cross-validation [CV] error; lower row: λ corresponding to the most regularized model within one standard error of the minimum) on the predictive accuracy of leave-one-out cross-validation (LOO-CV) clocks for *Odocoileus*: Individuals of both sexes from two species were used to build the LOO-CV clocks: mule deer (N = 50) and white-tailed deer (N = 33). The solid blue line represents the regression across both species, whereas the black dotted and grey dashed lines represent the mule deer and white-tailed deer regression lines, respectively. Predictive accuracy of LOO-CV clocks was evaluated using the median absolute error (MAE) and Pearson's correlation coefficient (indicated above each panel). The dashed line represents the diagonal (x = y).

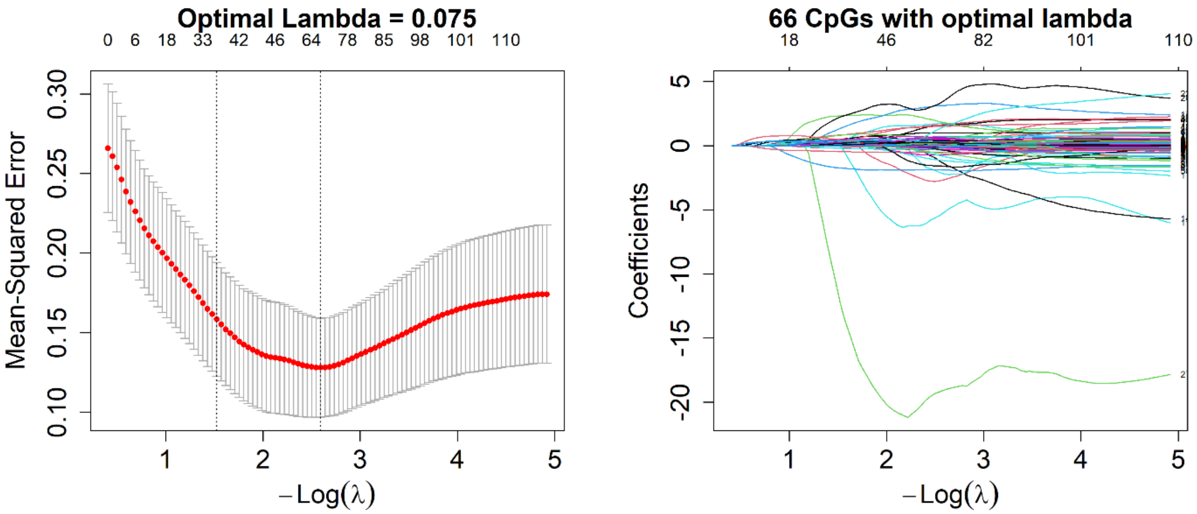


**Figure S4.** Selection of the optima regularization parameter (λ) and visualization of the regularization paths across λ values for the *Odocoileus* deer 10-fold cross-validated elastic-net model (α = 0.5), using log-transformed tooth age. <u>Left panel</u>: cross-validation curve used to select the optimal λ. Each point (red dot) represents the mean squared CV error, with error bars indicating ± SD. The vertical dotted lines represent the λ minimizing the CV error (lambda.min; here regarded as optimal λ here) and λ corresponding to the most regularized model within one standard error of the minimum (lambda.1se). <u>Right panel</u>: regularization path of CpG coefficients across λ values. Each curve corresponds to a CpG site (N = 27,270 CpGs), and CpGs with non-zero coefficients at the optimal λ were selected to fit the final *Odocoileus* deer elastic-net model.

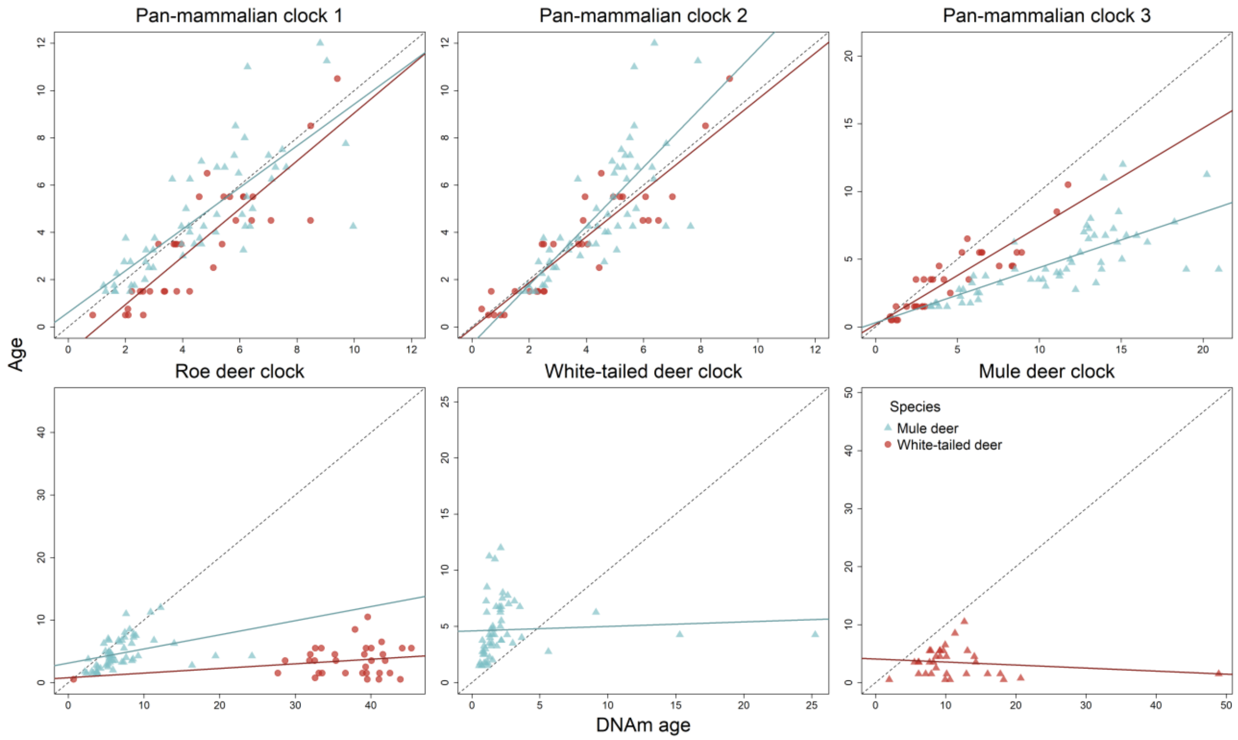


**Figure S5.** Prediction of mule deer and white-tailed deer chronological (tooth) age from methylation data using published epigenetic clocks and the mule deer clock developed in this study. The solid light-blue and red lines represent fitted regression lines for mule deer and white-tailed deer, respectively. Predictive accuracies of epigenetic clocks were: pan-mammalian clock 1 $r = 0.793$, MAE = 0.880 (mule deer: $r = 0.778$, MAE = 0.766; white-tailed deer: $r = 0.867$, MAE = 1.117), pan-mammalian clock 2 $r = 0.825$, MAE = 0.732 (mule deer: $r = 0.777$, MAE = 0.738; white-tailed deer: $r = 0.909$, MAE = 0.649), pan-mammalian clock 3 $r = 0.741$, MAE = 3.138 (mule deer: $r = 0.739$, MAE = 6.235; white-tailed deer: $r = 0.903$, MAE = 0.960), roe deer clock $r = -0.116$, MAE = 3.363 (mule deer: $r = 0.358$, MAE = 1.697; white-tailed deer: $r = 0.239$, MAE = 34.534), white-tailed deer clock $r = 0.060$, MAE = 2.806 (mule deer only), mule deer clock $r = -0.170$, MAE = 4.472 (white-tailed deer only).

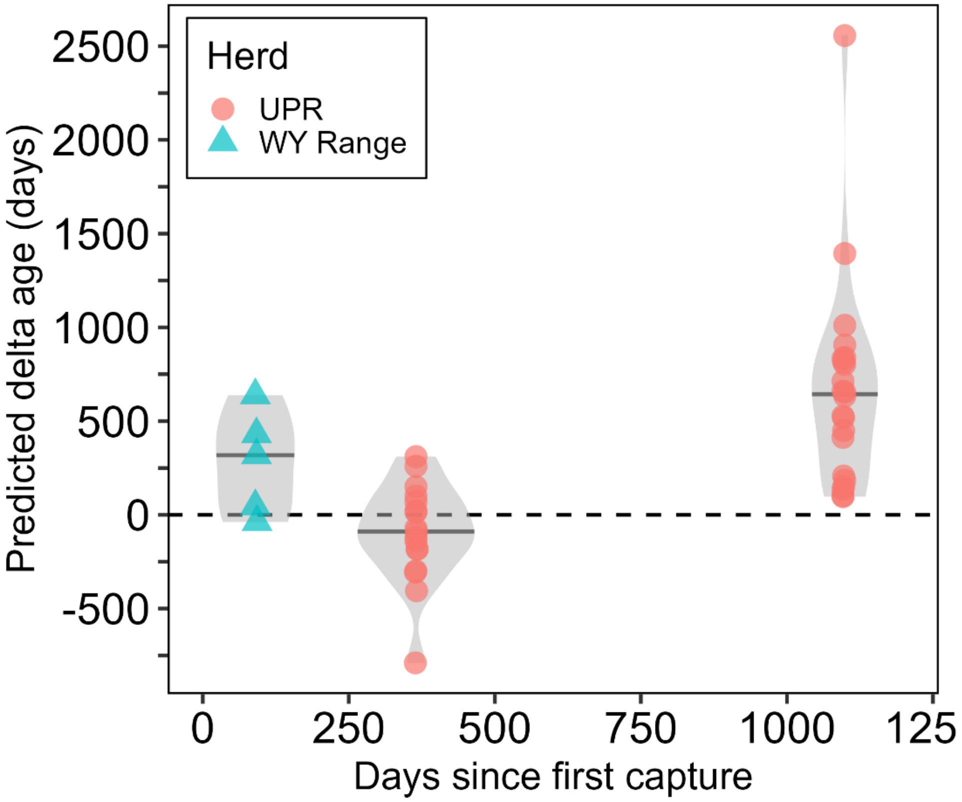


**Figure S6.** Difference in predicted DNA methylation (DNAm) age ($\Delta_{age}$) for 44 mule deer sampled twice in Wyoming, 2018-2022. Horizontal lines within the violin plots represent the median $\Delta_{age}$ for the three sampling intervals (90-93 days since $t_0$, 364-367 days since $t_0$, and 1096-1099 days since $t_0$).

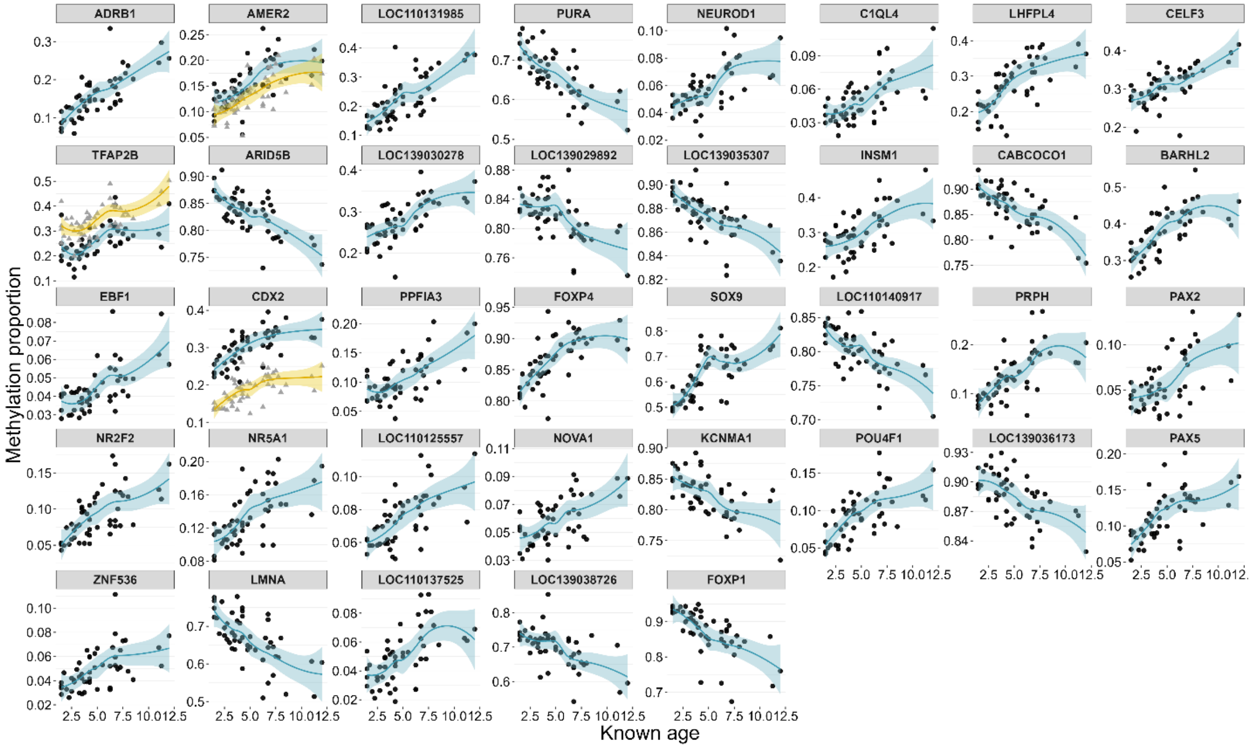


**Figure S7.** DNA methylation trajectories with age across 40 CpGs identified as associated with chronological age in the EWAS after Bonferroni correction for mule deer samples from Wyoming, 2018-2022. The name of the genomic feature or nearby genes is indicated above of each panel. Data points represent the methylation proportion of each individual included in the EWAS (N = 50). Lines represent LOESS fits, with shaded areas indicating the associated SE. Three genes were each associated with two distinct age-associated CpGs; to distinguish the trajectories of the two CpGs associated with each gene, data points for the second CpG are shown in grey, with its LOESS fit and associated SE in yellow. Panels are ordered by the EWAS P-value for the association with chronological age (from smallest to largest).
